# Common neural mechanisms control attention and working memory

**DOI:** 10.1101/2022.07.06.498935

**Authors:** Ying Zhou, Clayton E. Curtis, Kartik Sreenivasan, Daryl Fougnie

**Affiliations:** Program in Psychology, New York University Abu Dhabi, Abu Dhabi, UAE; Department of Psychology, New York University, New York, NY 10003; Center for Neural Science, New York University, New York, NY 10003; Program in Biology, New York University Abu Dhabi, Abu Dhabi, UAE

**Keywords:** attention, working memory, selection, fMRI, decoding

## Abstract

Although previous studies point to qualitative similarities between working memory (WM) and attention, the degree to which these two constructs rely on shared neural mechanisms remains unknown. Focusing on one such potentially shared mechanism, we tested the hypothesis that selecting an item within WM utilizes similar neural mechanisms as selecting a visible item via a shift of attention. We used fMRI and machine learning to decode both the selection among items visually available and the selection among items stored in WM in human subjects (both sexes). Patterns of activity in visual, parietal, and to a lesser extent frontal cortex predicted the locations of the selected items. Critically, these patterns were strikingly interchangeable; classifiers trained on data during attentional selection predicted selection from WM, and classifiers trained on data during selection from memory predicted attentional selection. Using models of voxel receptive fields, we visualized topographic population activity that revealed gain enhancements at the locations of the externally and internally selected items. Our results suggest that selecting among perceived items and selecting among items in WM share a common mechanism. This common mechanism, analogous to a shift of spatial attention, controls the relative gains of neural populations that encode behaviorally relevant information.

**Significance statement:** How we allocate our attention to external stimuli that we see and to internal representations of stimuli stored in memory might rely on a common mechanism. Supporting this hypothesis, we demonstrated that not only could patterns of human brain activity predict which items were selected during perception and memory, but that these patterns were interchangeable during external and internal selection. Additionally, this generalized selection mechanism operates by changes in the gains of the neural populations both encoding attended sensory representations and storing relevant memory representations.

## Introduction

Although theories of attention and working memory (WM) often emphasize their interrelatedness (Awh & Jonides, 2001; Chun, 2011; Cowan, 1998a; Gazzaley & Nobre, 2012; Oberauer, 2002), they are typically studied in isolation. Several lines of empirical evidence highlight commonalities between attention and WM. For example, attention and WM share similar capacity and resource limitations (Cowan, 1998b; Marois & Ivanoff, 2005; but see Fougnie & Marois, 2006) and engage similar brain regions (Awh & Jonides, 2001; Ikkai & Curtis, 2011; Jerde et al., 2012; LaBar et al., 1999; Pollmann & von Cramon, 2000; Ranganath & D’Esposito, 2005). Moreover, the process of rehearsing items (Awh et al., 1999, 2000; Awh & Jonides, 2001; Jha, 2002; Katus et al., 2014; Postle et al., 2004; Shen et al., 2015; Souza et al., 2020; Theeuwes et al., 2005) and suppressing distracting information (Gazzaley et al., 2005; Sreenivasan & Jha, 2007) in WM may be attention-based. In turn, representations in WM can guide how we attend to sensory information (Bahle et al., 2020; Gayet et al., 2013; Olivers et al., 2006; Sasin & Fougnie, 2020; Williams et al., 2022; Woodman & Luck, 2007). However, since attention and WM are complex processes involving multiple cognitive operations, it remains unclear which underlying components may be shared between the two. Here we focus on the process of selection, which refers to how task-relevant information is prioritized over task-irrelevant information in both attention and WM. Critically, selection helps mitigate the strict resource or capacity limitations of attention and WM (Marois & Ivanoff, 2005).

The notion that a common process both selects among external sensory information and among internal WM representations is intriguing because it appeals to an intuitive common mechanism used to highlight relevant information. Nonetheless, there is little evidence linking attentional selection and WM selection. Researchers have intensively investigated attentional selection using pre-cueing paradigms, where a cue indicates which forthcoming stimulus to attend (Eriksen & Yeh, 1985; Murphy & Eriksen, 1987; Posner, 1980). Behaviorally, pre-cueing benefits the processing of selected information (Carrasco et al., 2000; Pestilli et al., 2009) at the cost of the processing of unselected information (Pestilli et al., 2007; Pestilli & Carrasco, 2005). Both neurons (Bisley & Goldberg, 2003; Bushnell et al., 1981; Luck et al., 1997; Reynolds et al., 2000) and voxels (Gandhi et al., 1999; Hopfinger et al., 2001; Ikkai & Curtis, 2008; Liu et al., 2005; Serences & Boynton, 2007; Silver et al., 2007) with receptive fields that match the locations of pre-cued items exhibit increased activity relative to those that match the locations of unattended items. Theories of selective attention, including computational models, posit that the benefits of attention stem from gain enhancements within the populations of neurons encoding selected task-relevant stimuli (Carrasco, 2011; Reynolds & Heeger, 2009). To study the process of selection in WM, researchers have used retro-cueing experimental paradigms, where a cue presented after WM items have been encoded signifies which memorandum will later be tested (Griffin & Nobre, 2003). Behaviorally, the quality of memory is better for the retroactively cued item compared to non-cued items (Griffin & Nobre, 2003; Landman et al., 2003; Li et al., 2021; Souza et al., 2016). Neurally, WM representations are enhanced within the neural populations encoding the selected WM item (Ester et al., 2018; Lepsien et al., 2011; Sprague et al., 2016; Yoo et al., 2022).

We directly tested the hypothesis that a common neural mechanism underlies attentional and WM selection using fMRI and machine learning. Remarkably, classifiers trained on each type of selection were interchangeable in predicting the other, providing novel quantitative evidence for theories that posit a shared mechanism (Chun et al., 2011). In addition, using models of voxel receptive fields to visualize population activity, we observed elevated responses corresponding to the locations of the externally and internally selected items, suggesting that the shared selection mechanism involves differential gain.

## Materials and Methods

### Participants

Eleven neurologically healthy participants (ages 23-53; 6 females) with normal or corrected-to-normal vision participated in this experiment. The sample size was determined using previous fMRI studies comparing selection on perceptual and WM representations (Nobre et al., 2004; Tamber-Rosenau et al., 2011), and is equal to or larger than previous studies that compared within- and across-condition decoding performance of classifiers trained on fMRI data (Jerde et al., 2012; Kwak & Curtis, 2022; Rademaker et al., 2019), as well as those that used population receptive field-weighted reconstruction analysis (Kwak & Curtis, 2022; Yoo et al., 2022). Participants provided written informed consent in accordance with procedures approved by the Institutional Review Board at New York University.

### Experimental design

We generated stimuli and interfaced with the MRI scanner, button box, and eye tracker using MATLAB software (The MathWorks) and Psychophysics Toolbox 3 (Brainard, 1997). Stimuli were presented using a PROPixx DLP LED projector (VPixx) located outside the scanner room and projected through a waveguide and onto a translucent screen located at the head of the scanner bore. Participants viewed the screen at a total viewing distance of 63 cm through a mirror attached to the head coil. The display was a circular aperture with an approximately 32-dva (degrees of visual angle) diameter. A trigger pulse from the scanner synchronized the onsets of stimulus presentation and image acquisition.

Participants performed a pre-cue task and a retro-cue task in the two scanning sessions. The task procedures are illustrated in Figure 1A. The fixation symbol in both tasks was a centrally-presented filled circle with a 0.3-dva radius. Subjects were required to maintain fixation in the center of the screen. Each pre-cue trial began with a 750ms colored central fixation (0.4-dva radius) with three black placeholders. The color of the fixation indicated the target location in the upcoming stimulus screen. The distance from the screen center to the center of each placeholder was 6 dva, and the diameter of each placeholder was 8 dva. The pre-cue was followed by a 1500ms ISI, then by the stimulus for 1500ms. Stimulus presentation consisted of three Gabor patches, one in each placeholder. The three placeholders were in three different colors, and subjects had to select the target Gabor in the placeholder with the pre-cued color. The three colors used in each trial were randomly selected from four colors (RGB = [255, 0, 0], [0, 200, 0], [0, 0, 255], [255, 165, 0]) and randomly distributed across the three locations (Left, Right, Bottom) so that the target location could not be predicted by the representations of the pre-cue. The stimulus presentation was followed by a 750ms mask to diminish iconic memory (Sperling, 1960), then by a 3000ms delay. This was followed by the presentation of a probe, which consisted of a circle and an oriented line. The length of the line and the diameter of the circle were both 6 dva. Subjects had to judge if the line was rotated clockwise or counterclockwise compared to the orientation of the selected Gabor. We adjusted the difference between the orientations of target and probe to titrate the behavioral performance to approximately 80%. Specifically, the difference between the orientations of the probe and target started at 20° and either increased by 1° after each error trial or decreased by 1° after four continuous correct trials (cf. Levitt, 1971). Subjects responded by pressing ‘1’ for clockwise or ‘2’ for counterclockwise. The probe screen lasted for 2250ms regardless of subjects’ responses. Subjects then received feedback consisting of the selected Gabor overlaid with the probe. The color of the probe indicated whether the response was correct (green: correct; red: incorrect). The intertrial interval lasted for 9750ms. Each retro-cue trial began with a 1500ms stimulus screen, which contained three Gabor patches in three black placeholders. The stimulus was followed by a 750ms mask and a 1500ms ISI. Following the ISI, subjects saw a retro-cue consisting of three colored placeholders surrounding a colored fixation. The color of the fixation point matched the color of one of the placeholders and indicated the location of the target Gabor. The colors on each trial were randomly selected from the four possible colors (see above) and randomly distributed across the three locations. The delay, probe, feedback, and intertrial interval were the same as those in the pre-cue task. Each subject completed two scanning sessions consisting of 10 runs each, with five pre-cue runs and five retro-cue runs presented in an interleaved order. Each run contained 18 trials, yielding 180 trials per condition (90 per session). Each run started with 13 dummy TRs (9750ms) of a central fixation screen to allow for magnetic field stabilization.

**Figure 1.**
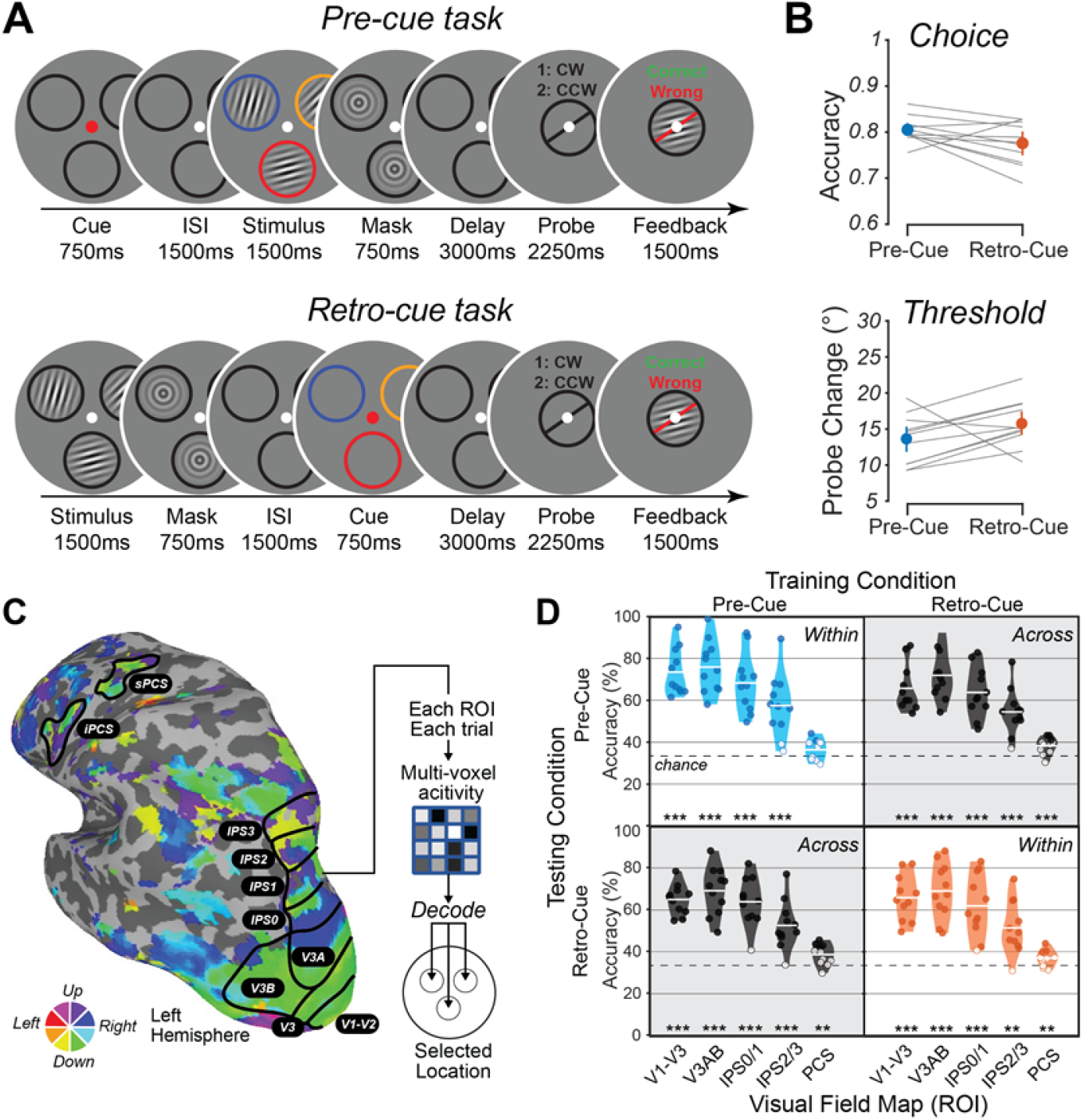
Methods, behavior, decoding analysis, and decoding results. **(A)** Pre-cue task: Fixation cued the task-relevant color. Then, three oriented Gabors appeared within a colored placeholder ring. Participants encoded the gabor whose ring color matched fixation. After a mask and delay, participants reported whether the orientation of the probe was tilted clockwise or counterclockwise to the memorized target orientation. Retroc-cue task: Participants encoded the orientations of three Gabors that were followed by a mask and short ISI. Then a retro-cue indicated the task-relevant color. Participants were instructed to remember the orientation of the item at the location matching the fixation color and later compare to the probe orientation. **(B)** Probe task accuracy and probe change magnitude in pre-cue (blue) and retro-cue conditions (red). Gray lines indicate individual subjects. The error bars denote 95% bootstrap CI. **(C)** Retinotopy for a representative subject. Color depicts voxels’ preferred polar angle projected to the cortical surface of the left hemisphere. **(D)** Decoding performance across ROIs, for within-condition decoding of pre-cued location (upper left), retro-cued location (lower right), and cross-condition decoding of location (gray backgrounds). Gray dashed line indicates chance accuracy (33.3%) (colored: individual performance was significantly higher than chance, p < .05; one-sided, FDR corrected across all ROIs for each decoding condition independently). The jittered dots indicate individual performance -filled dots indicate participants from whom decoding performance was higher than chance (p < .05, one-sided) and unfilled dots indicate participants with at-chance decoding performance. The horizontal line in the violin indicates the mean of accuracy. The stars in the bottom of each plot indicate the group-level performance is significantly higher than chance (***, p < .001; ** p < .01; * p < .05; one-sided, FDR corrected across all ROIs for each decoding condition independently).

### Eye movements

Eye position coordinates (x,y) and pupil size were recorded at 500Hz in the scanner using an EyeLink 2k (SR Research, Ontario, Canada). Prior to each scanning session, eye position was calibrated using a 9-point calibration. Eye data was preprocessed in Matlab using the freely available iEye toolbox (github.com/clayspacelab/iEye_ts) using the following steps. (1) Data was transformed from raw pixel screen coordinates into dva. (2) Extreme values associated with loss of track and blinks were removed. (3) Data was smoothed with a Gaussian kernel (5 ms SD). (4) Each trial was drift-corrected by taking the mean over known epochs when the participant was fixating (the whole delay in pre-cue task, and the first delay before the cue in the retro-cue task) and subtracting that value from the entire trial.

### fMRI Methods

#### MRI data acquisition

Participants underwent three fMRI scanning sessions - two for the main selection experiment and one for retinotopic mapping. All data were acquired at the NYU Center for Brain Imaging on a 3T Siemens Prisma Scanner with a 64-channel head/neck coil. Functional scans for the selection experiment were acquired using an EPI pulse sequence with 44 slices and a voxel size of 2.5^3^ mm (4x simultaneous-multi-slice acceleration; FoV = 200 × 200 mm, no in-plane acceleration, TE/TR = 30/750 ms, flip angle = 50°, bandwidth = 2290 Hz/pixel, 0.56 ms echo spacing, P → A phase encoding). Data for the retinotopic mapping session was acquired in a separate session at a higher resolution, with 56 slices and a voxel size of 2.0^3^ mm (4x simultaneous multi-slice acceleration, FoV = 208 × 208 mm, no in-plane acceleration, TE/TR = 36/1200 ms, flip angle = 66°, bandwidth = 2604 Hz/pixel, 0.51 ms echo spacing, P → A phase encoding). To correct for local spatial distortions in the functional data offline, we estimated a field map of the field inhomogeneities by acquiring pairs of spin echo images with normal and reversed phase-encoding directions with an identical slice prescription to the functional data and no simultaneous-multi-slice acceleration (TE/TR = 45.6/3537ms, 3 volumes per phase encoding direction). To enable precise localization of functional data, we collected T1-weighted whole-brain anatomical scans using a MP-RAGE sequence with 192 slices and a voxel size of 0.8^3^ mm (FoV = 256 × 240 mm, TE/TR = 2.24/2400ms, flip angle = 8°, bandwidth = 210 Hz/Pixel) for each participant.

#### MRI data preprocessing

T1-weighted anatomical images were segmented and cortical surfaces were constructed using the recon_all command in Freesurfer (version 6.0). Functional data were preprocessed with custom scripts using functions provided by AFNI. First we applied the *B*_0_ field map correction and reverse-polarity phase-encoding (reverse blip) correction. Next, we performed motion correction using a 6-parameter affine transform, aligned the functional data to the anatomical images, and projected the data to the cortical surface. Spatial smoothing (5 mm FWHM on the cortical surface) was applied to the retinotopic mapping data, but no explicit smoothing was applied to the data from the selection experiment. Data from the selection experiment was re-projected into volume space, which incurs a small amount of smoothing along vectors perpendicular to the cortical surface. Finally, we removed linear trends from the time series data, and then normalized (z-score) across all the time points within each run.

#### Estimating selection-related BOLD activity

To identify activity related to selection in both our tasks, we used the *3dDeconvolve* and *3dLSS* commands in AFNI (https://afni.nimh.nih.gov/) to implement a least-squares-separate general linear model (LSS-GLM) to the preprocessed blood-oxygen-level-dependent (BOLD) time series of the functional data. LSS-GLM has been shown to isolate single-trial activity in rapid event-related designs (Arco et al., 2018; Mumford et al., 2012). In the pre-cue condition, we modeled the selection, delay, and response events. The stimulus event was not modeled because it overlapped with the selection event. Based on previous estimates that symbolic central cues direct selection within 300ms (Carrasco, 2011), we set the duration of the selection event equal to one TR (750ms). The retro-cue model was similar to the pre-cue model, except the selection event was time-locked to the 750ms retro-cue screen. Each event was modeled as a boxcar with the duration of the event convolved with a hemodynamic response function (HRF; *GAM(10*.*9, 0*.*54)* in *3dDeconvolve*). The beta coefficients for the selection event in each trial were estimated separately, resulting in 180 GLM iterations for each task condition. In each iteration, we modeled the selection event on the trial of interest with a single regressor and the selection events on all other trials with a separate regressor. The delay and response events were modeled with one regressor each. Thus, the GLM for each task condition included 4 regressors on each iteration (selection for the trial of interest, selection for all other trials, delay, and response). In addition, the model included six regressors for head motion and four regressors for data drift. The procedure was repeated with each trial in turn serving as the trial of interest, resulting in 180 selection betas per condition. Selection betas were normalized via z-scoring on a voxel-by-voxel and run-by-run basis before further analysis.

### Population receptive field mapping

Population receptive field (pRF) mapping was conducted using the procedures described in (Mackey et al., 2017). Participants maintained central fixation while covertly tracking a bar aperture that swept across the screen in discrete steps in one of four orientations/directions: oriented vertically and traveling from left to right or right to left, or oriented horizontally and traveling from top to bottom or bottom to top. The bar aperture was divided into three rectangular segments (a central segment and two flanking segments) of equal sizes, each containing a random dot kinematogram (RDK). Participants’ task was to identify which of the two flanking RDKs had the same direction of motion as the central RDK. The dot motions of all the three segments changed with each discrete step. Participants reported their answer with a button press. We adjusted the coherence of the RDK using a staircase procedure to maintain accuracy at ∼75%. Each session contained eight to nine runs, each 5-minute run consisted of 12 sweeps, and each sweep consisted of 12 discrete steps (one step every 2s). The order of the four sweep directions was randomized within each run.

We fit the preprocessed BOLD time series for each voxel for each participant using the compressive spatial summation model (Kay et al., 2013)

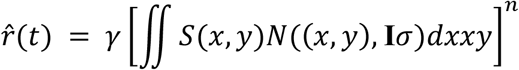

where *S* is a binary stimulus image (1 in where the stimulus was presented and 0 in where the stimulus was not presented), *N*((*x, y*), **I**σ) is a normal distribution with mean (*x, y*) and variance **I**σ^2^, where **I** is a two-dimensional identity matrix describing a circular, symmetric Gaussian. The parameters of this model are the voxel’s receptive field center (*x, y*) in dva, standard deviation σ in dva, amplitude γ, and compressive spatial summation factor *n* (where *n ≤* 1). Parameters were fit with a GPU-accelerated course grid search over parameters, followed by a local optimization method.

Voxels’ preferred phase angle (arctan 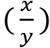) and eccentricity 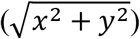 were visualized on the cortical surface. To define retinotopically organized regions of interest (ROIs), we restricted our analysis to voxels with greater than 10% response variability explained by the pRF model. We then drew ROIs on each participant’s cortical surface using reversals of the voxels’ preferred phase angle as boundaries between neighboring visual regions (Mackey et al., 2017). We defined four ROIs in anterior and dorsal visual areas (V1–V3AB), four ROIs along the caudal–rostral intraparietal sulcus (IPS0–IPS3), and two ROIs along the dorsal-ventral precentral sulcus region (sPCS and iPCS), each with a full visual field representation (see Figure 2A for an illustration of the ROIs for one subject). Further fMRI analyses were conducted in ROIs that combined regions which share a foveal confluence (V1, V2, and V3; IPS0 and IPS1; IPS2 and IPS3; Mackey et al., 2017; Wandell et al., 2005, 2007). We also combined voxels in sPCS and iPCS into a single PCS ROI in order to roughly match the size of our other ROIs, although the results were comparable for the individual PCS ROIs.

**Figure 2.**
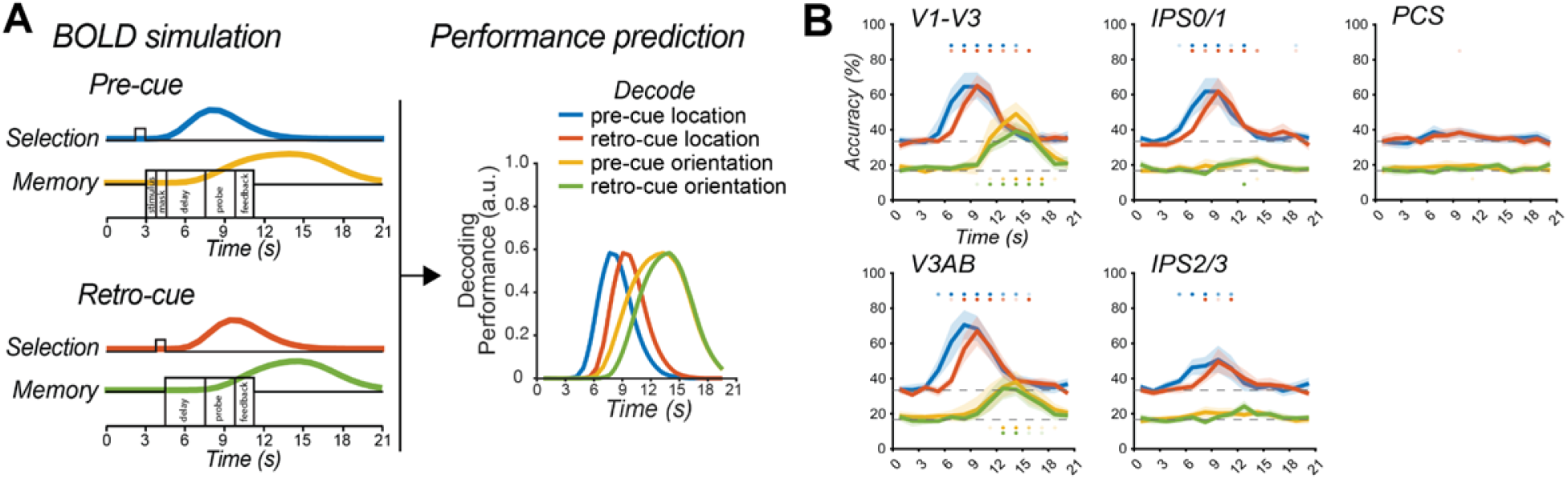
Temporal evolution of selection and memory decoding. **(A)** Simulating selection and memory related activity and predicting the performance of decoding target location and orientation. The neural activities for selection and memory were simulated by convolving the hypothetical duration with an HRF. The performance prediction was proportional to the amplitude of neural activities in arbitrary units. **(B)** The time-series of decoding performance for target location and orientation. The gray dashed lines key the chance for decoding location (33.3%) and orientation (16.7%). Blue and red lines stand for decoding location in pre-cue and retro-cue conditions, respectively. Yellow and green lines stand for decoding orientation in pre-cue and retro-cue conditions, respectively. The colored lines depict the mean of the performance for all subjects. The filled area around the line keys the 95% bootstrap CI. The dots on the top and bottom indicate the performance at the corresponding time-points is significantly higher than chance (the darkest color means p < .001; the medium color means p < .01; the lightest color means p < .05; one-sided, FDR corrected across all time points for each ROI and decoding condition independently).

### Decoding analyses

We examined whether the machine learning algorithm trained by selection-related activity in the pre-cue task can decode the internally selected location in the retro-cue task, and vice versa. To decode the selected location, we trained sparse multinomial logistic regression (SMLR) classifiers in Matlab using the Princeton MVPA toolbox (http://www.csbmb.princeton.edu/mvpa). The SMLR classifier is widely used to decode multi-class conditions (Krishnapuram et al., 2005; Pereira et al., 2009). The z-scored beta coefficients for the selection event in each trial estimated using the LSS-GLM were used to train the classifiers. First, the classifiers were tested on the dataset in the same task condition as training (i.e., within-condition decoding) to examine if the neural representations we extracted could predict the selected location (see an illustration in Figure 1C). Since there were three possible locations, we used a leave-three-trials-out cross-validation scheme, in which the classifier was trained on the data from all but three trials (one for each location) and then tested on the left-out trials in one task condition for each cross-validation fold, resulting in 60 cross-validation folds. Decoding accuracy was computed by comparing the true location labels with the classified labels across the 60 cross validations. Next, the classifier was trained on all of the data from one task condition and tested on each trial from the other task condition (i.e., across-condition decoding) to examine if the neural representations were comparable in the two task conditions. To increase the reliability of our decoding estimates, we repeated the entire procedure 10 times, taking the mean of the decoding accuracy across iterations as the final within- and across-condition decoding accuracy.

To estimate the BOLD activity for selection and memory phases in the task, we computed the time series of decoding performance. We segmented the z-scored preprocessed BOLD time series from 0 to 21s (i.e., the end of ITI) relative to the trial onset, and trained classifiers separately using the BOLD signal averaged across every 2TRs. One set of classifiers was trained to decode target location, signifying the selection process, while the other set of classifiers was trained to decode target orientation, signifying the maintenance process. Decoding of the target location was conducted using the leave-three-trials-out cross-validation scheme described above, while decoding of target orientation was carried out using a leave-six-trials-out cross-validation scheme, corresponding to the six possible Gabor orientations.

### Estimating population-level activity modulation

We tested whether external and internal selection both cause an activity increase in voxels whose receptive fields match the selected item’s location relative to the unselected items’ locations. To estimate the response of the neural populations at the selected location, we reconstructed a pRF-weighted map using the selection-related activity. This procedure essentially projects voxel activity in each ROI to visual space in screen coordinates (Kok & de Lange, 2014; Kwak & Curtis, 2022; Yoo et al., 2022). In each ROI, for every selected location (left, right, bottom) in each task condition (pre-cue, retro-cue), we created a reconstructed map using

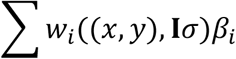

where *w*_*i*_((*x, y*), **I**σ) is the weight associated with the *i*th voxel at location (*x, y*), and *β*_*i*_ is the averaged GLM-acquired selection period *β* at voxel *i* in trials with the same selected location. We defined *w*_*i*_ as the receptive field of voxel *i*, which was a two-dimensional Gaussian with mean (*μ*_*x*_, *μ*_*y*_) and variance **I**σ^2^, where **I** is a two-dimensional identity matrix describing a circular, symmetric Gaussian (see Figure 3A for an illustration). For each pRF-weighted selection activity map, we calculated the mean of selection-related activity in the selected and non-selected locations, and averaged across three target location conditions. The activity difference between the selected and non-selected locations was taken as the activity modulation due to selection.

**Figure 3.**
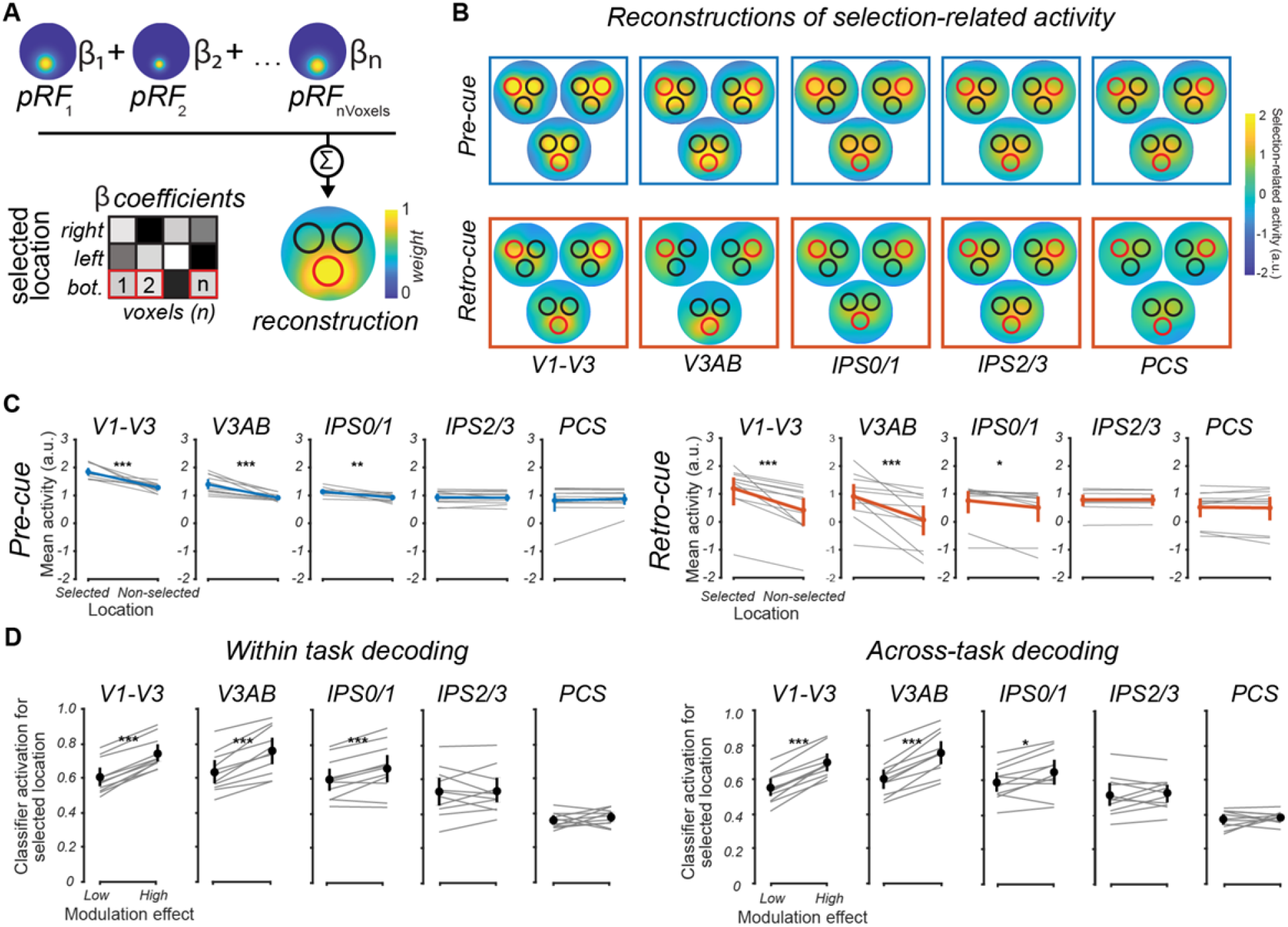
Reconstructing the topographic patterns of selection. **(A)** Procedure for reconstructing the pRF-weighted selection activity maps. For each condition and ROI, we multiplied the GLM-obtained selection-related β coefficients for each voxel, for each location (i.e., left, right, down) separately, by its pRF model parameters (i.e., position, size) and summed across all voxels. This procedure projects the relative population activity within the ROI in brain space into screen coordinates of visual space. **(B)** The pRF-weighted reconstruction maps for each condition and ROI. The circles in the map mark the three placeholder rings shown during the experiment, and the red ring keys the selected location. **(C)** The mean of pRF-weighted selection-related activity in the selected and non-selected locations in each task and ROI for individual subjects (gray) and across subjects (blue for pre-cue and red for retro-cue). These results indicate that changes in population gain underlie attentional and WM selection (see Figure 3-1 for simulation results that provide additional evidence in favor of a gain mechanism). **(D)** The relationship between the activity modulation observed in (C) and decoding results. Within-task (left) and across-task (right) decoding strength in the form of classifier activation is shown separately for trials with low and high modulation. Error bars are the 95% bootstrap CI. Stars in the top indicate the significance of the difference between the two conditions indicated in the x-axis (***, p < .001; ** p < .01; * p < .05; one-sided, FDR corrected across all ROIs for each task condition independently).

### Relationship between decoding results and population-level activity modulation

To investigate the relationship between our decoding results and population reconstructions, we conducted a median split of trials based on the magnitude of population-level modulation for the selected location and compared the relative classifier activation for the selected location for low- and high-modulation trials. Classifier activation for each spatial location was calculated as the sum over voxels of the classifier training weights assigned to each voxel multiplied by the BOLD activity on a given trial, and was normalized to sum to 1 over the three classes. We used the relative classifier activation (where higher activation can be taken as greater classifier evidence for that class) for the selected location as a measure of the strength of classifier evidence for each of the three possible locations.

### Distinguishing between possible mechanisms underlying enhanced population response

To explore whether observed increases in population response were consistent with relative gain modulation, we used a simulation to compare the expected influence on the spatiotopic population response under two plausible mechanisms - multiplicative gain (Herrmann et al., 2010; Reynolds & Heeger, 2009) and pRF shifts (Vo et al., 2017) - with our observed population-level modulations. Each voxel’s activity was simulated by the cumulative distribution function

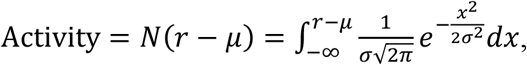

where *r* denotes the radius of the selected Gabor stimulus, *μ* denotes voxel’s estimated pRF center, and σ denotes voxel’s estimated pRF size (given by the SD of the best-fit Gaussian). In the multiplicative gain model, selection-related modulation, *m*, of activity for voxels with pRF centers within the selected Gabor was given by

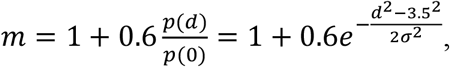

where *d* denotes the distance between voxel’s pRF center and the center of the selected Gabor and σ denotes voxel’s estimated pRF size; (cf. Figure 1 in Reynolds & Heeger (2009); Figure 5 in Herrmann et al. (2010)). In the pRF shift model, we assumed that voxels’ pRF centers would shift a distance of *s* towards the center of the selected location according to

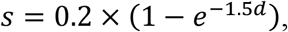

where *d* denotes the distance between voxel’s pRF center and the center of the selected Gabor (cf. Figure 2e in Vo et al. (2017)). To assess which model best described our data, we calculated the sum of squared error between simulated and observed activity modulation. The simulation process and result are shown in Figure 3-1.

### Statistical analysis

For our behavioral analyses, we used paired-sample *t*-tests to examine differences between performance across tasks. For our fMRI analyses, all statistics were calculated using a nonparametric permutation analysis (Rademaker et al., 2019). This method is appropriate here because there was no a *priori* reason to believe that the data would be normally distributed. Specifically, we repeated each analysis with shuffled trial labels (i.e., selected location: left, right, bottom) 1000 times to build an empirical null distribution of the test statistic of interest (e.g., decoding accuracy). For individual-level analyses, the percentage of the empirical null distribution that was equal to or larger than the real data was taken as the *p*-value. For group-level analyses, our test statistic was the *t*-value derived from a paired *t*-test of the real data (vs. zero or chance), and the empirical null distribution was the corresponding *t*-value for each of the 1000 iterations. The *p*-value was the percentage of the null distribution equal to or larger than the true *t*-score.

### Data Availability

Behavioral, eye-tracking, and MRI data, and analysis code are available at https://osf.io/jqu95/.

## Results

### Behavioral performance

The accuracy of memory judgments in the pre-cue and retro-cue tasks indicated that participants performed each task well (Figure 1B). Each condition utilized a separate staircase with a target accuracy of 80%. As expected, we found no significant difference in accuracy between the pre-cue (81%) and retro-cue (78%) conditions, *t*(10) = 2.084, *p* = .064, Cohen’s *d* = .628, *BF*_10_= 1.421. At threshold, the angular differences between the sample and probe orientations in the pre-cue (13.64°) and retro-cue conditions (15.76°) did not significantly differ, *t*(10) = 1.856, *p* = .093, Cohen’s *d* = .560, *BF*_10_= 1.074, suggesting that performance was well-matched across the two tasks.

### Decoding target location from isolated selection-related BOLD activity

First, we estimated selection-related activity on each trial using a GLM that modeled each of the trial components for each cueing condition (**Materials and Methods**). Second, for each condition separately we trained classifiers to predict the cued locations (i.e., left, right, down; Figure 1C). For both pre-cue and retro-cue trials, most of the ROIs could successfully decode the selected location (Figure 1D), confirming that our decoding procedures were robust. Third, to test our main hypothesis regarding a shared neural selection mechanism, we performed across-condition decoding. We trained classifiers using pre-cue data and attempted to decode the cued location using retro-cue data. Similarly, we trained classifiers using retro-cue data and decoded the cued locations using pre-cue data. Critically, we found that most of the ROIs could successfully decode the selected location across cueing conditions (Figure 1D). The performance of across-condition decoding was approximately 90% of within-condition decoding, suggesting that the neural activity patterns were nearly interchangeable across the two tasks. We ruled out that these results were due to gaze shifts during selection (Hedge & Leonards, 2013; Theeuwes et al., 2005, 2009); classifiers trained on eye position were unable to predict the selected location (all FDR corrected *p*s ≥ .05).

### Temporal evolution of selection and memory

To confirm that our decoding results indeed represented selection as opposed to incidental maintenance of the selected location, we examined the time course of decoding accuracy. This analysis was motivated by the fact that location information was essential during the selection process itself, but ceased to be task-relevant once selection was complete (as the memory probe was presented at fixation). Thus, we predicted that the time course of decoding accuracy for location would rise following the onset of the selection cue and return to chance once selection was complete, while the time course of orientation decoding (signifying the maintenance, retrieval, and response processes) would peak after selection and remain above chance for the duration of the trial. To formalize this prediction, we separately convolved the selection regressor and the combined delay + response + feedback regressors from our GLM with the HRF to simulate the expected time courses of the selection and memory processes, respectively (Figure 2A). We then compared these simulated time courses with the actual decoding time course for location and orientation. Critically, the decoding time course analysis was completely independent from the GLM used to derive the simulated time courses (see **Materials and Methods**). The decoding accuracy time series for location and orientation (Figure 2B) closely matched our simulated time courses. Specifically, the rapid rise and fall of location decoding in visual and parietal cortex matched the simulated time course of selection, whereas orientation decoding in visual cortex mirrored the simulated memory time course. Moreover, location decoding accuracy peaked earlier in the pre-cue relative to the retro-cue condition, consistent with the relative timing of selection in the two tasks. Together, these observations support our claim that location decoding reflected the selection process while orientation decoding reflected memory processes.

### Reconstructing maps of selection-related activity

Although these results derived from machine learning approaches provide strong evidence for interchangeable patterns of activity during attentional and WM selection, it is notoriously difficult to make direct inferences about neural mechanisms based on significant classification (Freeman et al., 2011; Naselaris et al., 2011; Serences & Saproo, 2012). We used each voxel’s pRF parameters (i.e., position and size) to project selection-related activity within each ROI to the screen coordinates of visual space (Figure 3A). We used these reconstructions to visualize how selection impacted the distribution of activity within the retinotopically organized maps. Notably, we found increased activity in the portions of the maps containing the selected target relative to the distractors for both the pre-cue and retro-cue conditions (Figure 3B). To quantify these results, we compared the mean activity within the selected and non-selected locations. For both pre-cue and retro-cue conditions, the mean activity at the selected location was higher in V1-V3, V3AB, and IPS0/1 (Figure 3C). To investigate whether this relative difference in activity explained the significant cross-decoding between pre-cued and retro-cued selection, we conducted a median split of trials for each subject based on the magnitude of population modulation for the selected location and compared classifier evidence (quantified as the classifier activation for the selected location relative to the activation for the non-selected locations) for low- and high-modulation trials. Both within- and across-task classifier evidence were significantly greater for high-modulation relative to low-modulation trials in V1-V3, V3AB, and IPS0/1 (Figure 3D). Such relative difference in spatiotopic activity modulation explains why we found significant cross-decoding between selection during pre- and retro-cue tasks. Most provocatively, these findings are highly consistent with the effects of multiplicative gain enhancement that have been observed at the population (Corbetta et al., 1990; Sprague & Serences, 2013) and single-neuron (Connor et al., 1997; McAdams & Maunsell, 1999; Treue & Martínez Trujillo, 1999) level; as such, our results point to a plausible neural mechanism underlying a shared mechanism of external and internal selection.

To provide further support for the notion that gain enhancement is the mechanism underlying our observed changes in population response, we used simulations to directly compare two mechanisms of selection. We simulated fMRI data expected under multiplicative gain and under pRF shifts and compared these data to our observed data in Figure 3C. Critically, while both multiplicative gain (Herrmann et al., 2010; Reynolds & Heeger, 2009) and RF shifts (Vo et al., 2017) have been observed with fMRI during spatial selection, these mechanisms make different predictions about the spatial pattern of modulation in voxels that represent the selected location. Specifically, the gain model predicts that the largest enhancement in response would occur in voxels whose RFs cover the center of the selected location, while the RF shift model predicts a larger enhancement in voxels whose RFs cover the periphery of the selected location (i.e., voxels whose RF centers shift from outside the selected location to inside the selected location). Our observed data was significantly more consistent with the gain model in V1-V3, V3AB, and IPS1/0 (all FDR corrected *p*s *≤* .004), but not in IPS2/3 or PCS where the two models did not significantly differ (all FDR corrected *p*s ≥ .080; Figure 3-1). These findings bolster the claim that gain modulation supports a common selection operation.

## Discussion

While both attention and WM involve the preferential processing of task relevant information, here we addressed the extent to which they draw on shared mechanisms. Importantly, the intuitive and appealing notion that attentional and WM selection reflect a single underlying process (Chun et al., 2011; Kiyonaga & Egner, 2013) lacks direct evidence. Behavioral studies using dual task paradigms report inconsistent effects of a concurrent attention-demanding task on WM selection (Janczyk & Berryhill, 2014; Lin et al., 2021); but see (Hollingworth & Maxcey-Richard, 2013; Makovski & Pertzov, 2015; Rerko et al., 2014). Furthermore, while neural studies consistently observe that attentional and WM selection evoke activity in overlapping brain regions (Gazzaley & Nobre, 2012; Griffin & Nobre, 2003; Kuo et al., 2009; Nobre et al., 2004), this is a *qualitative* and not a *quantitative* conclusion, and crucially ignores the possibility that selection processes may draw on distinct neural mechanisms that coexist in the same brain regions. As an illustrative example, perception of first-order and second-order motion have been found to activate nearly identical regions in V1 (Nishida et al., 2003), but a quantitative comparison demonstrated that different populations were responsible for each (Ashida et al., 2007). Our findings establish a stronger case for the overlap of selection operations for perceptual and mnemonic information than prior work by identifying interchangeable patterns of activity in human visual, parietal, and to a lesser extent frontal cortices during attentional and WM selection. An important distinction between our work and existing MVPA studies comparing attention, perception, and WM is that these other studies have focused on overlap in perceptual and mnemonic representations (Jerde et al., 2012; Serences et al., 2009), as opposed to the operations that facilitate the use of these representations to guide behavior. In addition, our results represent an advance over previous studies that found task-generalized decoding only in prefrontal cortex (Panichello & Buschman, 2021) or not at all (Tamber-Rosenau et al., 2011) by providing evidence for a generalized selection mechanism that spans multiple cortical regions. An important area for future studies will be to identify the specific contexts that engage this shared mechanism.

Another key theoretical advance of our work lies in identifying a putative mechanism – activity enhancement of spatiotopic population-level responses in the selected location (i.e., gain enhancement) – that underlies selection in attention and WM. Enhanced sensory responses during attentional selection is a well-established finding at the population level (Corbetta et al., 1990; Mangun et al., 1993; Sprague & Serences, 2013), most likely due to a multiplicative scaling of neuronal responses within the attended receptive field (Connor et al., 1997; McAdams & Maunsell, 1999; Treue & Martínez Trujillo, 1999; Williford & Maunsell, 2006) that drives preferential processing of perceptual information. In contrast, there is comparatively little evidence for the role of gain enhancement in WM selection besides qualitative similarities between WM-induced modulations of visual processing and the enhancement of sensory processing during attentional selection (Awh et al., 2000; Merrikhi et al., 2017; Sreenivasan et al., 2007; Sreenivasan & Jha, 2007). Our findings suggest that the vast literature on multiplicative gain can be leveraged to better understand how we select information from within WM. According to theory, increases in neural gain enhance the signal-to-noise ratio, and therefore precision, of neural representations (Ma et al., 2006; Zemel et al., 1998). Thus, the gain increases we observed associated with selection might control which items are prioritized in WM. Notably, while gain enhancement is the most plausible mechanism based on our simulation results (Figure 3-1), we do not claim to have conclusively ruled out all mechanisms of selection. Importantly, our results constrain theory by demonstrating that any plausible mechanism would need to produce equivalent relative activity modulation at the populational level.

Do our decoding results reflect the selection process itself or the outcome of the selection process (i.e., the consequence of having selected a particular location; Myers et al., 2017)? We considered this question in two ways – first by examining the time course of location decoding to distinguish between selection and memory for the selected information. We found that the strongest decoding of location was time-locked to the selection events in both tasks, with decoding accuracy quickly falling to chance once the location information was no longer relevant. In contrast, orientation decoding peaked later after selection and remained above chance for the entire memory delay. This pattern of findings indicates a transient process by which the task-relevant location was selected followed by prolonged maintenance of the target’s orientation. Thus, location decoding in our data likely represents the selection process itself. The time-limited representation of selection that we observed may help explain important discrepancies in the behavioral literature – some studies find dual-task costs between attention and selection in WM (Janczyk & Berryhill, 2014; Lin et al., 2021) and others do not (Hollingworth & Maxcey-Richard, 2013; Makovski & Pertzov, 2015; Rerko et al., 2014). We argue that studies that have failed to observe interference generally assume that attention is continuously applied to maintain selection in WM (Hollingworth & Maxcey-Richard, 2013), while those that find interference generally put the secondary task temporally near the selection cue (Janczyk & Berryhill, 2014). Further elucidation of the temporal profiles of selection in WM should be an important area for future investigation.

Second, despite the fact that our study was not specifically designed to distinguish the sources controlling selection from the effects of selection, our findings intriguingly point to potentially distinct roles of visual and association cortex. In visual cortex, not only could we decode the selected location, but in later time points we could decode the target orientation held in memory, suggesting a role in WM storage (Curtis & D’Esposito, 2003; Serences et al., 2009; Sreenivasan et al., 2014), potentially as a consequence of receiving top-down selection signals (Rahmati et al., 2018; Sprague et al., 2016; Yoo et al., 2022). On the other hand, in parietal and frontal cortex, we had robust decoding of selected location, while decoding of memorized orientation was inconsistent across the delay and across ROIs (Figure 2B), consistent with the idea that gain enhancement in topographically organized regions of parietal and frontal cortex reflects the sources of top-down signals controlling which locations are selected. While we cannot completely rule out the possibility that unsuccessful orientation decoding was due to larger RF sizes or increased spatial heterogeneity in the representation of features in these regions (see Ester et al. (2015) for a successful demonstration of orientation decoding in frontoparietal cortices), our findings are reminiscent of the role of frontoparietal cortex in orienting attention and prioritization (Corbetta & Shulman, 2002; Jerde et al., 2012; Serences & Yantis, 2006) and gating (Chatham et al., 2014; Frank et al., 2001), and spatial cognition more broadly (Corbetta et al., 2000; Heilman et al., 1985; Mackey et al., 2016; Mesulam, 1999; Srimal & Curtis, 2008; Szczepanski et al., 2010; Vandenberghe et al., 2001; Yantis et al., 2002). It is worth noting that a previous study that failed to find a common multivoxel pattern across attentional and WM selection used different features (location and object) for selection across tasks (Tamber-Rosenau et al., 2011). Given that neural substrates of selection are sensitive to differences in the medium of selection (Giesbrecht et al., 2003), conclusions about a common selection mechanism ought to be drawn from comparisons which rigorously match the attention and WM tasks. Here we compare across the same selection medium (location), equating relevant features, as well as behavioral performance. Consistent with the idea that WM selection relies on internally-directed shifts of attention that highlight task-relevant information, our results suggest that a common mechanism underlies selection during attention and WM.

## Acknowledgements

We thank New York University’s Center for Brain Imaging for technical support. This research was supported by the United States National Eye Institute (NEI) (R01 EY-016407 to C.E.C. and R01 EY-027925 to C.E.C.), the Abu Dhabi Department of Education and Knowledge (ADEK) Abu Dhabi Award for Research Excellence (AARE) (AARE19-230) to K.K.S, and the Research Enhancement Fund (RE177) from New York University Abu Dhabi to D.F.

## Extended data

**Figure 3-1.**
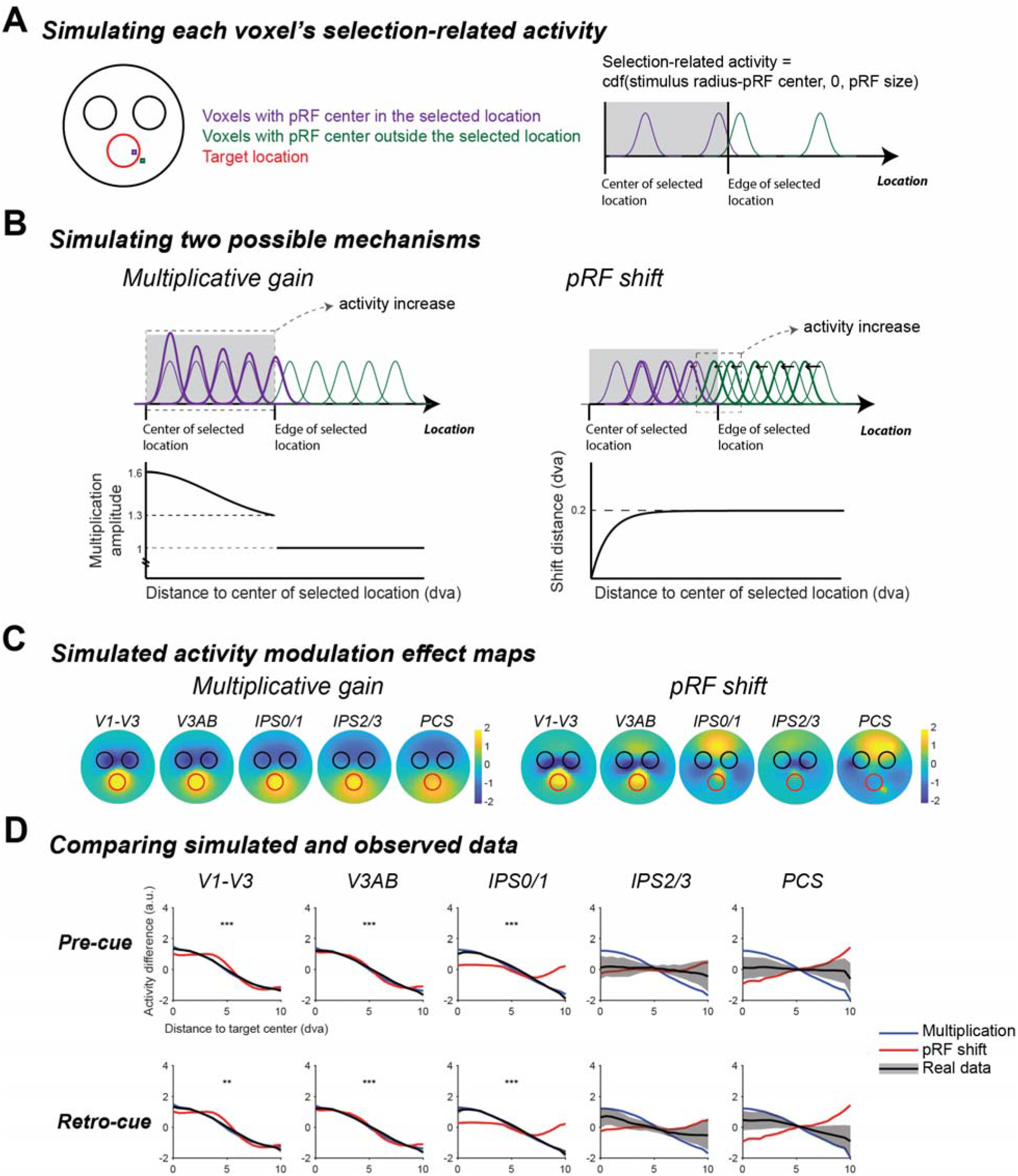
Simulating multiplicative gain and pRF shift mechanisms. **(A)** Schematic illustrating the simulation of selection-related activity. **(B)** Schematic illustrating the simulation of the two possible mechanisms. In the multiplicative gain model, activity enhancement was modulated by multiplying the selection-related activity of voxels with pRF centers in the selected location. In the pRF shift model, voxels’ pRF centers were shifted towards the center of the selected location. **(C)** The simulated activity modulation maps for the two possible mechanisms. We reconstructed the pRF-weighted map using the simulated selection-related activity using the same procedure as that used to generate Figure 3B, isolated activity modulation effect by subtracting the mean of the pRF weighted map for the two non-selected locations from that for the selected location (e.g., middle – (left+right)/2), and collapsed the modulation maps across three selected locations. The circles in the map mark the three placeholder rings shown during the experiment, and the red ring keys the selected location. **(D)** Comparison between simulated and observed data. The modulation effect decays as voxel’s pRF center gets away from the selected location. The blue and red lines are for multiplicative gain and pRF shift mechanism respectively, and the black line is for the observed data. The filled area around the line indicates the 95% bootstrap CI. The stars in the top of each plot indicate that the sum of squared error between the multiplication simulation data and real data was significantly lower than that between the pRF shift simulation data and real data (***, p < .001; ** p < .01; * p < .05; one-sided, FDR corrected across all ROIs). The observed data was significantly more consistent with the multiplicative gain model in most of our ROIs.

