## Supplementary figures and images for "Common neural mechanisms control attention and working memory"

### Figure 3-1

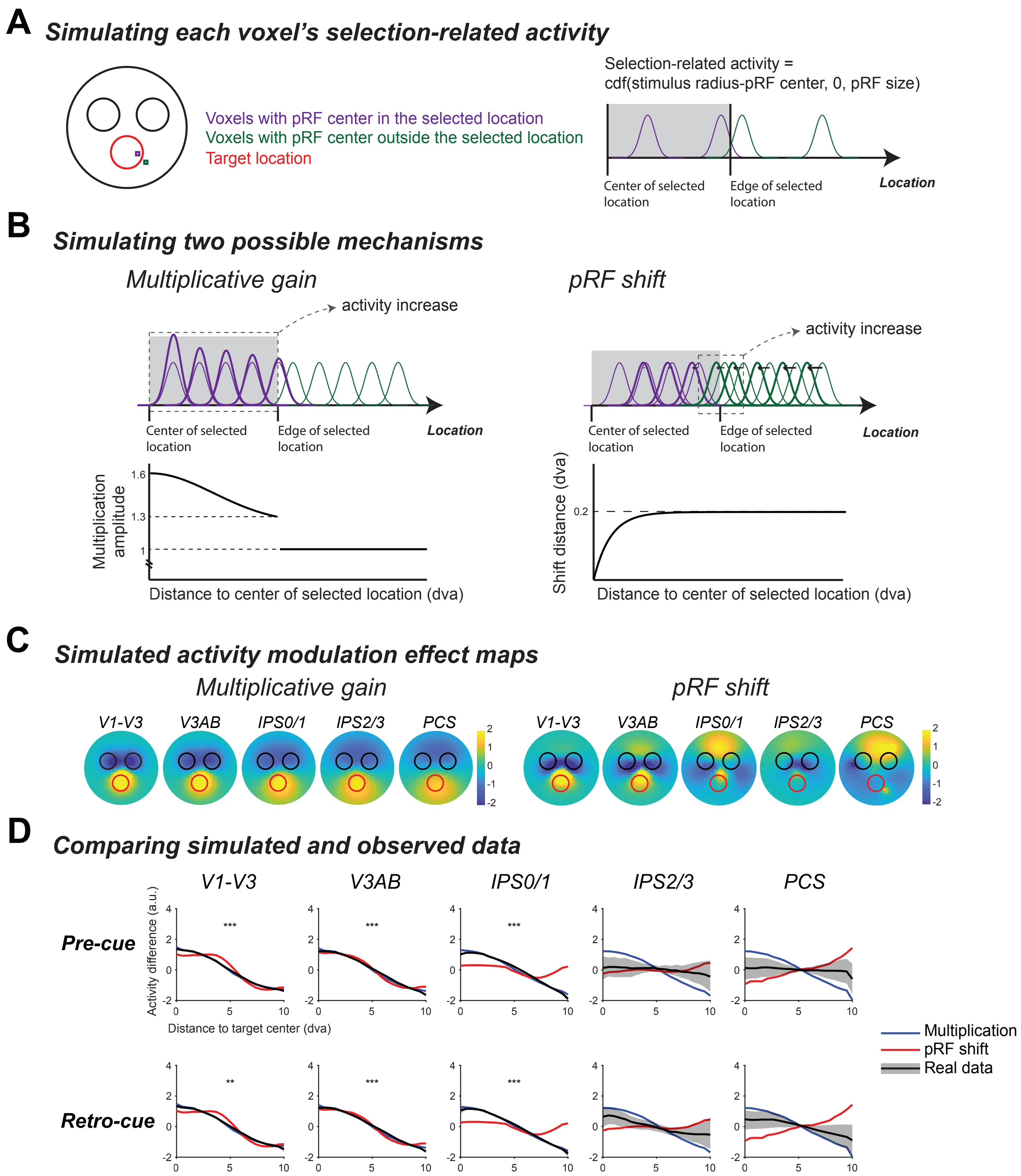
